# Kinship and genetic variation in aquarium-spawned *Acropora hyacinthus* corals

**DOI:** 10.1101/2022.10.18.512587

**Authors:** Elora H. López-Nandam, Cheyenne Y. Payne, J. Charles Delbeek, Freeland Dunker, Lana Krol, Lisa Larkin, Kylie Lev, Richard Ross, Ryan Schaeffer, Steven Yong, Rebecca Albright

## Abstract

Recent scientific advances in *ex situ* system design and operation make it possible to complete gametogenic cycles of broadcast spawning corals. Breeding corals in aquaria are critical advances for population management, particularly genetic rescue and assisted gene flow efforts. Genetic rescue projects for corals are already underway to bring threatened species into *ex situ* culture and propagation, thereby preserving standing genetic variation. However, while breeding corals is increasingly feasible, the consequences of the aquarium environment on the genetic and phenotypic composition of coral populations is not yet known. The aquarium environment may in itself be a selective pressure on corals, but it also presents relaxed selective pressure in other respects. In 2019 and 2020, gravid *Acropora hyacinthus* coral colonies were collected from Palauan reefs and shipped to the California Academy of Sciences (CAS) in San Francisco. In both years, gametes were batch-fertilized to produce larvae that were then settled and reared to recruits. As of April 2021, when they were sampled for sequencing, 23 corals produced at CAS in 2019 and 16 corals produced at CAS in 2020 had survived for two years and one year, respectively. We sequenced the full genomes of the 39 offspring corals and their 15 potential parents to a median 26x depth of coverage. We find clear differential parentage, with some parents producing the vast majority of offspring, while the majority of parents produced no surviving offspring. After scanning 12.9 million single nucleotide polymorphisms (SNPs), we found 887 SNPs that may be under selection in the aquarium environment, and we identified the genes and pathways these SNPs may affect. We present recommendations for preserving standing genetic variation in aquarium-bred corals based on the results of this pilot project.

## INTRODUCTION

Breeding animals and reintroducing their offspring to the wild as a means of bolstering a threatened population started in the 1960’s, with the successful breeding and reintroduction of the Arabian oryx (Spalton et al., 1999). By the 1980’s, a major goal of conservation breeding programs was not simply to increase the number of animals in a population or species, but to maintain genetic variation in populations (Ballou, 1984). As DNA sequencing technology improved and became less expensive, some conservation breeding programs began to incorporate genetic analyses to determine kinship among their animals, and to prevent or reduce inbreeding by selecting unrelated individuals to mate with one another (Fienieg and Galbusera, 2013).

Conservation breeding programs at zoos have now successfully bred and reintroduced several species that are threatened in the wild. The most famous of these programs have been focused on terrestrial megafauna, primarily mammals and birds, e.g. California condors (Chemnick et al., 2000) and Florida panthers (Johnson et al., 2010). Aquatic conservation breeding programs have led to reintroductions of freshwater amphibians, molluscs, and fish, e.g. hellbender salamanders (Ettling et al., 2017), Oregon spotted frogs (Howell et al., 2021), freshwater mussels (Araujo et al., 2015), and desert pupfish (Koike et al., 2008). Conservation breeding initiatives for marine animals have traditionally focused on fish hatcheries (Fisch et al., 2015) and aquaculture of invertebrates like tridacnid clams (Frias-Torres, 2017).

The decline of many marine invertebrate species, and in particular corals, highlights an urgent need for breeding programs of corals for conservation and restoration (van Oppen et al., 2015; Humanes et al., 2021). Increasing threats to coral reefs globally have sparked a need for new, scalable conservation and management solutions. The majority of coral nursery and propagation techniques are currently based on fragmentation and asexual propagation of coral clones (Henry et al., 2021). While these methods can increase coral cover in a particular region, they have the potential to decrease the genetic variation within the population because the fragments are genetic clones of each other. Standing genetic variation, which is comprised of all unfixed alleles in a population, can contribute to adaptation when a new or heightened selective pressure changes the frequency of one or more alleles in a population (Hermisson and Pennings, 2005; Barrett and Schluter, 2008). The greater the genetic variation present in the population, the greater the likelihood that one of those variants may be adaptive in the future (Hoffmann and Willi, 2008; Eizaguirre and Baltazar-Soares, 2014). Standing genetic diversity allows for adaptation through weakly adaptive alleles that exist in the population at low frequency but become more advantageous in the presence of a new or heightened selective pressure. As selective pressures intensify, adaptive alleles increase the likelihood of an individual’s survival and become more common in the population as the organisms without the adaptive allele die and/or the adapted organisms reproduce more successfully (Hoffmann and Willi, 2008). Thus, as water temperatures and acidity rise in coral habitats globally, the capacity for coral populations to adapt to those changes will depend in large part on having sufficient genetic variation, such that some of these variants may confer a selective advantage to environmental change (Bay et al., 2017).

The importance of standing genetic variation for species adaptation is a major reason why conservation breeding programs seek to preserve as much standing genetic variation as possible. However, animals in a zoo, aquarium, or hatchery are exposed to different selective pressures than they would experience in their natural habitat (Frankham, 2008). While it is certainly true that the aquarium environment eliminates many potential selective pressures, such as predation, it may inadvertently introduce others. In addition to unintentional selection, there are other challenges that conservation breeding programs encounter, including reduced genetic variation due to inbreeding, which occurs when highly related organisms produce offspring, and genetic drift, the stochastic fixation of alleles that can have large effects when the population size is small (Chargé et al., 2014). Some simulations have suggested that the reduction of fitness due to loss of adaptive variation in zoo or aquarium-bred animals will result in lower population census within generations once they are reintroduced to the wild (Willoughby and Christie, 2019). Empirical studies on fisheries have shown that fish born in hatcheries have lower fitness than their wild counterparts after just a single generation (Kostow, 2004; Araki et al., 2007; Christie et al., 2012; Wakiya et al., 2022). While some studies have documented changes in fitness or selection in zoo, aquarium, or hatchery-bred populations, none have explored the biological pathways or functions that are under selection in these settings. Combating loss of fitness in conservation breeding programs will necessitate strategies that minimize the frequency of detrimental alleles and maximize retainment of adaptive and neutral alleles. In addition, conservation breeding may affect the holobiont, or the full suite of microorganisms that live in and on the host animals of interest. Some tridacnid clam conservation breeding programs incorporate measurement of the symbiotic zooxanthellae that the juvenile clams uptake (Niartiningsih et al., 2020). This aspect of conservation breeding is still underexplored, but is likely to be critical for animals like corals and clams for which symbiosis with algae and other microbes is critical for survival.

Recent advances in long-distance transport of corals, as well as improvements in system design to mimic seasonal water temperature fluctuations, solar irradiance, lunar cycles, and diel cycles *ex situ*, have allowed predictable coral spawning *ex situ* to become possible (Craggs et al., 2017, 2018; O’Neil et al., 2021). Public aquariums are an ideal setting for coral spawning and breeding pilot programs, as they have the infrastructure, resources, and technical expertise in the form of personnel who know how to keep corals alive and healthy. Pairing aquarium expertise in animal husbandry with next-generation sequencing allows for new insights into how breeding corals *ex situ* affects the genetic composition of aquarium-born animals, and the extent to which the aquarium environment introduces novel selection pressures. The Coral Spawning Lab at California Academy of Sciences is a collaborative endeavor between two departments within the museum: the Steinhart Aquarium and the Institute for Biodiversity and Sustainability Sciences. In 2019 and 2020, gravid corals were imported from Palau and spawned in the lab. The gametes were collected and batch-fertilized, and the aquarium-bred offspring were reared to juvenile coral colonies. In 2021, we sequenced the full genomes of the corals that spawned in 2019 and 2020 (F0 generation), as well as the offspring (F1 generation) that survived to be two years old (2019 F1s) and one year old (2020 F1s) at the time of tissue sampling. We show evidence of differential parentage, with some parents producing many F1s that lived to be at least one year old and many parents producing none. We also identify single nucleotide polymorphism (SNP) candidates that may have been under selection due to the lab environment in both 2019 and 2020, and highlight the functional pathways that these SNPs may affect. These data serve as a first indicator of how breeding corals *ex situ* may influence the genetic and phenotypic composition of the resulting aquarium-born population. Based on these data, we provide recommendations for minimizing inbreeding, genetic drift, and selection for the aquarium environment in aquarium-bred corals.

## MATERIALS AND METHODS

### Coral Collection

Gravid *Acropora hyacinthus* coral colonies were collected in Palau (Bureau of Marine Resources permit number RE-19-07 and CITES permit PW19-018) in February 2019 and February 2020, in anticipation of the 2019 and 2020 spawns, respectively. Colonies were transported to the Coral Spawning Lab at the California Academy of Sciences, where they were kept on a Palauan cycle (lighting and temperature) until spawning, with methods adapted from Craggs et al. (2017). See Supplementary Methods for seasonal temperature settings and lighting regimes for 2019, 2020, and 2021.

### Gamete collection, fertilization, larval rearing, and settlement

Colonies were monitored for spawning activity on nights 0 – 15 after the simulated full moons (0-15 nights after full moon, referred to as NAFM) in March 2019 and April and May 2020. Spawning occurred on 6-9 NAFM (March 27-30) in 2019 and 12-15 NAFM (April 19-22) and 10-12 NAFM (May 17-19) in 2020 (Table 1). Following release, gamete bundles were collected in 50 ml falcon tubes, labeled, and set aside for fertilization. Tubes were gently agitated to assist disaggregation of gamete bundles to release of eggs and sperm. Bundles disaggregated ~60 min after release whereupon eggs and sperm from each colony were combined in 0.45 μm filtered seawater (FSW) and left to batch fertilize for 60 min. Following fertilization, embryos were rinsed in 0.45 μm FSW and gently transferred to continual flow larval cones maintained at 26-27 °C in 0.45 μm FSW. Cultures were maintained at the recommended rearing density of ~1 larva mL^−1^ (Pollock et al., 2017) over the course of 4-7 days until competent to settle. Once competent, larvae were settled onto pre-conditioned (>4 months) aragonite tiles (Ocean Wonders ®), inoculated with symbionts isolated from parent colonies, and reared for 1-2 years. The 2019 F1s were reared in the Coral Spawning Lab (CSL) at the California Academy of Sciences. Due to COVID-associated closures, the 2020 F1s were reared in an offsite lab to enable daily access and care during critical early life stages. Descriptions of both aquarium setups are provided below. For both systems, herbivores (urchins, snails, fishes) were included in the tanks with corals to help minimize algal growth.

**Table 1.**
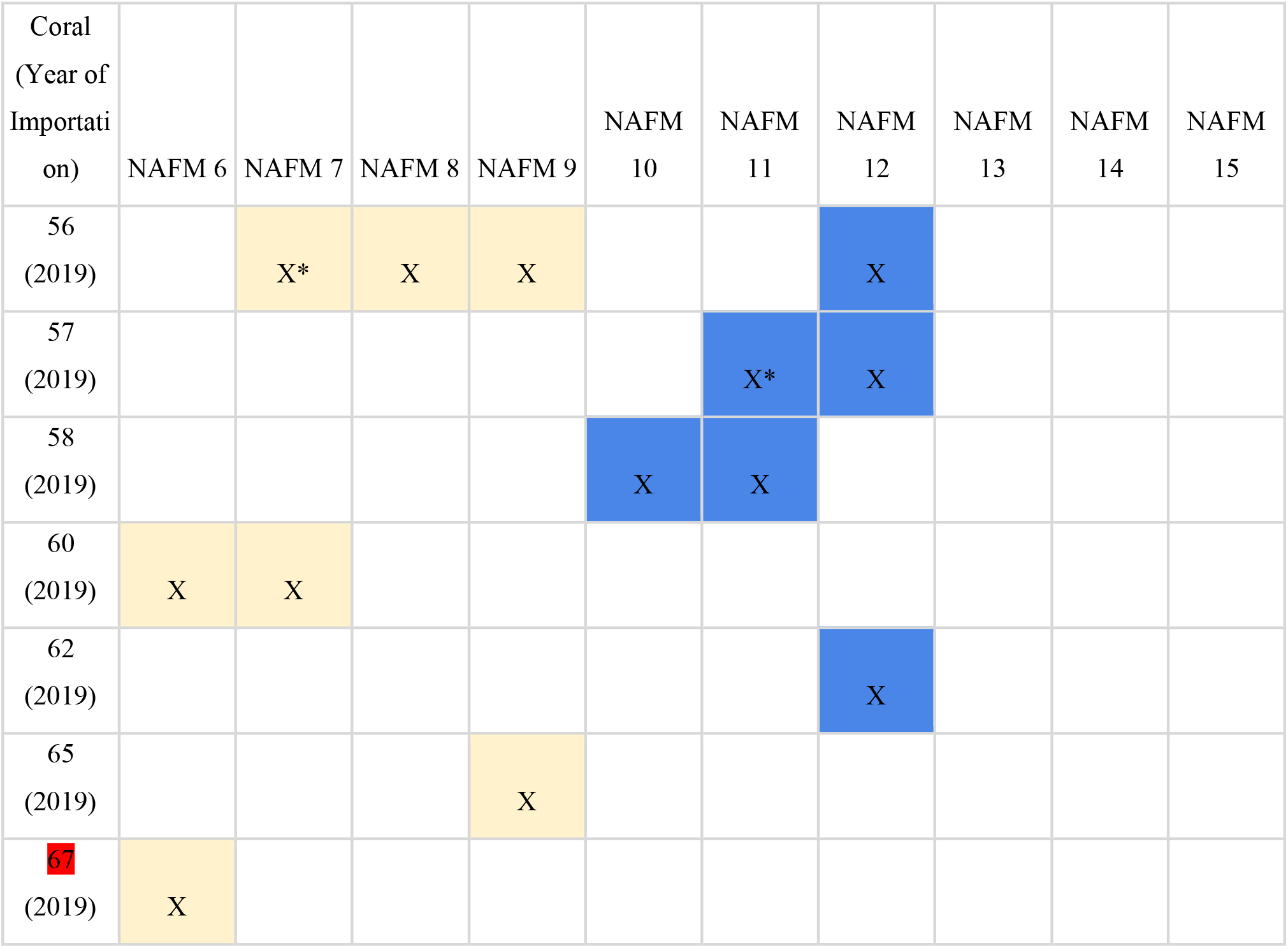
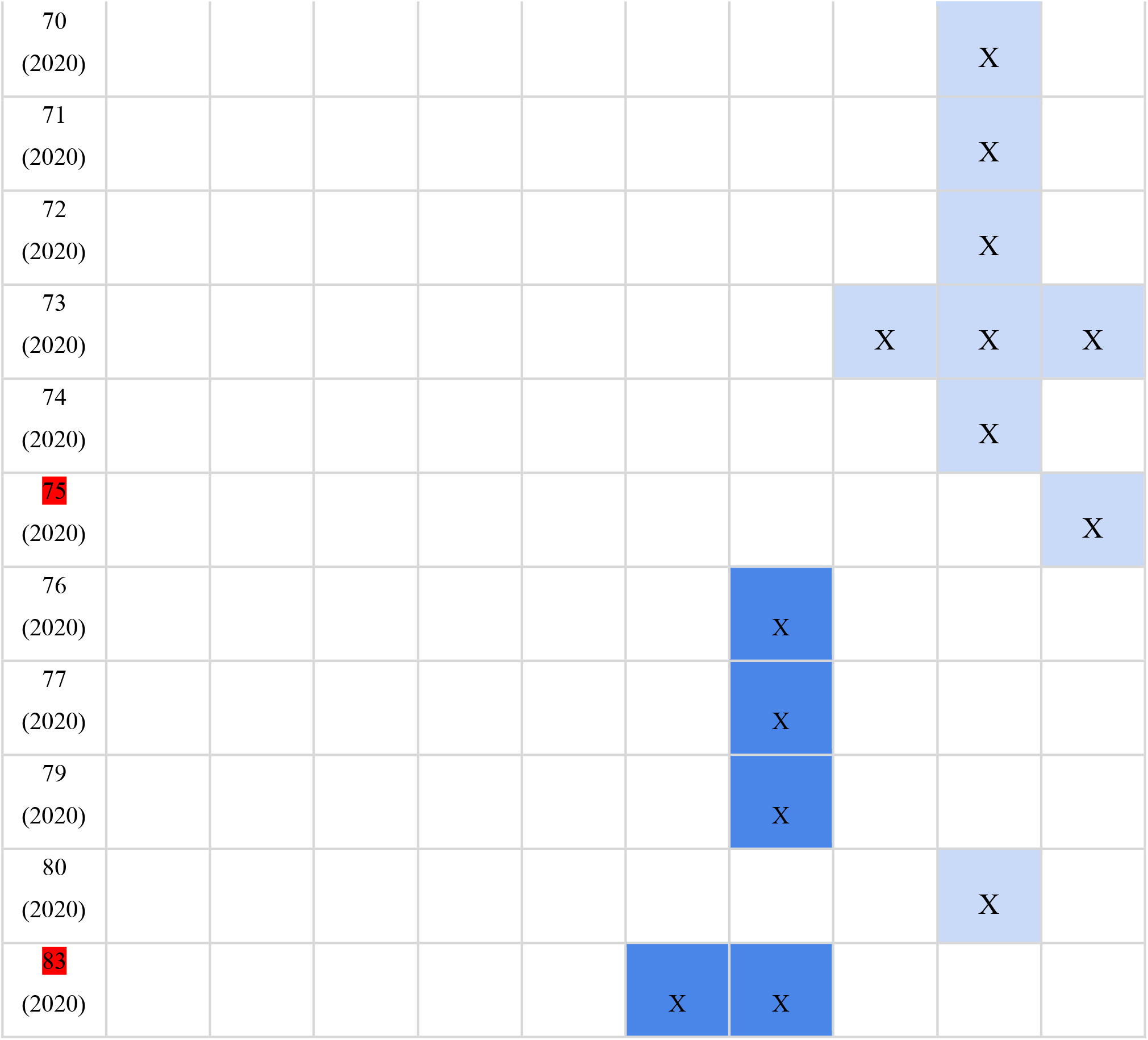
Spawning activity relative to the night after full moon (NAFM), where X indicates that a given colony spawned on a given night. Yellow is March 2019, light blue is April 2020, dark blue is May 2020. Spawners that were not sequenced are highlighted in red. Asterisks represent days on which colonies dribbled, or released just a few bundles.

### Aquarium setup (CSL)

The Coral Spawning Lab (CSL) at the California Academy of Sciences was built in 2018 and is nested in the Academy’s Steinhart Aquarium. The laboratory aquarium system is in a dark room, consisting of an outer vestibule for a two-step entrance that protects the tank area from light pollution, and uses temperature and lighting control to manipulate coral spawning, with modifications from Craggs et al. (2017). The CSL is a closed 438-gallon (1658 L) saltwater system consisting of six 60-gallon (227 L) (36”x30”x14”) tanks and a filtration system. Parameters for the system were programmed to mimic the water temperature and light cycles of Palau (Supplementary Methods). Temperatures ranged from 26-28 °C (79-83 °F), and each tank was lit by two Ecotech Marine Radion XR15 G4 lights, with water motion provided by 2 Neptune Systems WAV pumps per tank. The system uses artificial seawater with the following parameters: nitrate (NO_3_^−^) 4.3 mg/l, phosphate (PO_4_^3-^) 0.05 mg/l, salinity 33-36 parts per thousand (ppt), pH 8-8.4, alkalinity 2.6-3.0 mEq/L, magnesium 1400 mg/L, and calcium 380-430 mg/L (see Supplementary Methods for full artificial seawater recipe). Corals were fed once a day with a rotating mixture of live phytoplankton, live *Artemia nauplii*, copepods, rotifers (Live S-Rotifers, Reed Mariculture, Campbell, California, USA), and particulate reef food for filter feeders (BeneReef Reef Food, Benepets, Salt Lake City, Utah, USA). The systems are controlled by Neptune® Systems Apex controllers which automate temperature and light cycles to mimic seasonal changes in seawater temperature and lighting.

### Offsite aquarium setup

The offsite aquarium used to rear the 2020 F1s consists of a central aquarium system originally established in 2003, and a satellite life support system that was established in 2006 and upgraded for coral spawning in 2018. The total system volume is approximately 450 gallons (1703 L). Two 67-gallon (254 L) tanks (34”x 24”x18”) were used to rear out the 2020 coral recruits. These two tanks had a turnover rate from the life support system of approximately five times per day. The system uses filtered natural salt water at a temperature of 25.5 °C (78F), pH 8.1-8.3, nitrate (NO_3_^−^) 50 mg/l, phosphate (PO_4_^3-^) 0.9 mg/l, salinity 33-36 parts per thousand (ppt), alkalinity 3.0 mEq/L, magnesium 1250 mg/L, and calcium 400 mg/L. Each larval tank was lit by a single Ecotech Marine Radion gen 4 XR30 (and later Neptune systems SKY light), with water motion provided by 2 Neptune Systems WAV pumps per tank. The systems are controlled by Neptune Systems Apex controllers.

### Sample collection, preparation, and sequencing

In April 2021, one 1 cm branch was broken off of each adult spawner and stored in a 1.5 mL tube of 99% ethanol. Three of the adult spawners (CA56, CA60, CA65) had been sampled and sequenced to high depth of coverage in 2019 (López-Nandam et al. 2022), so they were not resampled at this time (Supplementary Table 1). For the 2019 F1s, smaller branch clips of 5-8 polyps were taken and for the 2020 F1s, 2-4 polyps were scraped into ethanol. Samples were stored at 4 °C until extraction.

Two polyps were scraped off of each ethanol-preserved sample for each DNA extraction. Three of the spawning colonies (CA72, CA74, and CA80) died prior to tissue sampling; therefore, preserved sperm from the 2020 spawn was used for DNA extraction instead of adult polyp tissue of these colonies. Three adult spawners (CA67, CA75, and CA83) were not sequenced because they died prior to sampling and no sperm was preserved from them. DNA was extracted from sampled tissue using Qiagen DNEasy kits following the Blood and Tissue protocol with modifications specifically for genomic DNA extraction from corals (Baums and Kitchen 2020) and a few further modifications (Supplementary Methods). Extracted DNA was sent to Texas A&M Agrilife Bioinformatics and Genomics Service (College Station, TX, USA) for whole-genome library preparation using a NEXTFLEX Rapid XP DNA-Seq Kit HT (PerkinElmer, Waltham, MA, USA). Libraries were sequenced at the Texas A&M Agrilife Bioinformatics and Genomics Service on one NovaSeq 6000 S4 lane at 26.1 ± 5.8 depth of coverage per sample across the full genome (2 s.d.) (Illumina, San Diego, CA, USA). Three of the parent libraries-CA56, CA60, and CA65-were sequenced previously at Chan-Zuckerberg Biohub (San Francisco, CA, USA) and one of the parent libraries-CA74-was sequenced previously for genome assembly at Dovetail Genomics (Scotts Valley, CA, USA) (López-Nandam et al., 2022).

### Read Mapping and SNP calling

Adapters were trimmed from reads using trimmomatic, version 0.39 (Bolger et al., 2014). Trimmed reads were mapped to the *Acropora hyacinthus* v1 reference genome (López-Nandam et al., 2022) using BWA version 0.7.17-r1188 with the bwa-mem algorithm (Li and Durbin, 2009). Duplicate reads were removed with Picardtools MarkDuplicates version 2.25.7. Depth of coverage across the genome for each sample was calculated using Genome Analysis Toolkit Version 4.2.0.0 DepthofCoverage tool (Van der Auwera et al., 2013). Haplotype calling was performed with the Genome Analysis Toolkit version 4.2.0.0 Haplotypecaller tool. Following GATK’s best practices for variant calling, we combined GVCFs from the same coral colony into a multi-sample GVCF using CombineGVCFs. Joint genotype calling was then performed on each multi-sample GVCF using GenotypeGVCFs with the option –all-sites to produce genotypes for both variant and nonvariant sites. The genotype-called multi-sample VCFs were filtered to remove all loci where one or more samples were missing a genotype call, and then further filtered so that depth of coverage was greater than 10 for every sample and GQ was greater than 30 at any sample using BCFtools (Danecek et al., 2021). We filtered for biallelic single nucleotide polymorphisms (SNPs) using BCFtools. The finalized, filtered VCF was annotated with snpEff (Cingolani et al., 2012) configured with the *Acropora hyacinthus* v1 genome (López-Nandam et al., 2022). All sequenced samples are listed with number of mapped and unmapped reads per sample in Supplementary Table 1. For the complete read mapping and SNP calling pipeline, including full commands with all parameters, see https://github.com/eloralopez/AquariumBreedingGenomics.

### Parentage Analysis

To calculate identity-by-descent (IBD) between individuals, we used Plink v2.00a2.3LM (Chang et al., 2015) to generate a pi_hat score of the proportion of sites in IBD for every pair of individuals in the population, including all wild-sourced spawners and all offspring produced in the aquarium. Using plink2, we made pairwise comparisons among every pair of individuals across the full population. For each pair, we found how many loci were state 0 (no shared alleles), state 1 (one shared allele), or state 2 (two shared alleles), and the proportion of alleles estimated to be in identity by descent, pi_hat = P(IBD=2) + 0.5*P(IBD=1) (Supplementary Table 2). Two clones would yield a pi_hat score of ~1, full siblings and parent-offspring pairs have a pi_hat of ~0.5, and half siblings have a pi_hat of ~0.25. To identify parent-offspring pairs, we filtered offspring-spawner pairs for pi_hatscores approximately equal to 0.5. We then checked putative parent-parent-offspring trios against the spawning date matrix (Table 1) to determine whether putative parent pairs spawned on the same day. All identified trios were validated as viable based on the spawning date matrix. Two offspring, CA2019-24 and CA2020-10, each had a PI_HAT of ~0.5 with just one of the sequenced parent colonies. Using the spawning date matrix (Table 1), we determined that the second parent for each of these offspring was one of the two spawners that were not sequenced (CA67 was the second parent of CA2019-24, CA58 was the second parent of CA2020-10).

### F_ST_ Outlier Identification

To identify SNPs with outlier F_ST_ values, we calculated F_ST_ between spawners and offspring for each year using VCFtools --weir-fst-pop version 0.1.16 (Danecek et al., 2011). We used a custom R script to identify SNPs with F_ST_ values in the 99th percentile in each cohort, and then identified the SNPs that were in the 99th percentile for both cohorts. We further filtered this subset to only include SNPs for which allele frequency, calculated in plink2 with the --freq option, changed in the same direction from F0 to F1 in both cohorts, i.e. the allele frequency increased in both cohorts or decreased from both cohorts. We refer to these SNPs as the “shared outliers” (Supplementary Table 3). See https://github.com/eloralopez/AquariumBreedingGenomics for scripts.

### Allele and Genotype Frequencies

In addition to calculating allele frequencies in full cohorts, we also calculated allele frequencies in subsets of the cohorts to compare observed allele frequencies with those expected under Mendelian inheritance. We calculated allele frequencies in the 22 full siblings produced in 2019, the 6 full siblings produced in 2020, and the two sets of respective parents for these offspring in plink2 with the --freq option (Supplementary Tables 4-5).

To compare observed genotype frequencies to those expected under Hardy-Weinberg equilibrium, we calculated genotype frequencies in plink2 with the --geno-counts option. Autosomal Hardy-Weinberg equilibrium exact test statistics were calculated in plink2 with the -- hardy option (Supplementary Table 6). For custom R and python scripts used in these analyses see https://github.com/eloralopez/AquariumBreedingGenomics.

### GO enrichment of outliers

To determine whether specific biological pathways were enriched in the set of shared outliers, we performed a Gene Ontology (GO) term enrichment analysis. We pulled the *Acropora hyacinthus* transcripts that either overlapped outlier coordinates or that were closest to an outlier in a noncoding region. Given that there are currently no functional annotations or mapped GO terms for predicted *A. hyacinthus* transcripts, we performed a nucleotide BLAST search of the outlier-associated *A. hyacinthus* transcript sequences against a local database of starlet sea anemone (*Nematostella vectensis*) cDNA sequences downloaded from EnsemblMetazoa (genome assembly ASM20922v1; see command line used in Appendix 1). *N. vectensis* is the closest cnidarian species supported in the extended Ensembl database with annotated GO terms. For each transcript, we chose the top *N. vectensis* BLAST hit by selecting the hit with the highest bit score and with the lowest e-value, which measure sequence similarity and the number of expected hits with the same quality by chance, respectively (see Appendix 1). We used the corresponding *N. vectensis* transcript IDs as the target subset for GO enrichment.

We generated a gene universe with all 20,4681 *N. vectensis* genes with annotated GO terms, and matched Ensembl transcript IDs with GO terms using the R packages “biomaRt” v2.50.3 (Durinck et al., 2009) and “GSEABase” v1.56.0 (Morgan et al., 2022). We note that the *N. vectensis* gene set is not perfectly representative of the *A. hyacinthus* gene set; however, we expect the majority of the genome to be conserved between these species. We used the GSEAbase implementation of a hypergeometric test to test for overrepresented GO biological pathways, molecular functions, and cellular compartments in the set of outlier-associated genes.

## RESULTS

### 2019 and 2020 spawns in the aquarium

Four of fourteen *Acropora hyacinthus* colonies that were imported in 2019 spawned in March 2019. Eleven of thirteen colonies that were imported in 2020 spawned in 2020 - seven in April and four in May. Additionally, four of the colonies that were imported in 2019 spawned in 2020, for a total of 15 spawners in 2020 (eleven 2020 imports and four 2019 imports). Spawning activity for all 2019 and 2020 corals is presented in Table 1. By April 2021, there were 23 surviving offspring from the 2019 spawn and 16 surviving offspring from the 2020 spawn (Figure 1).

**Figure 1.**
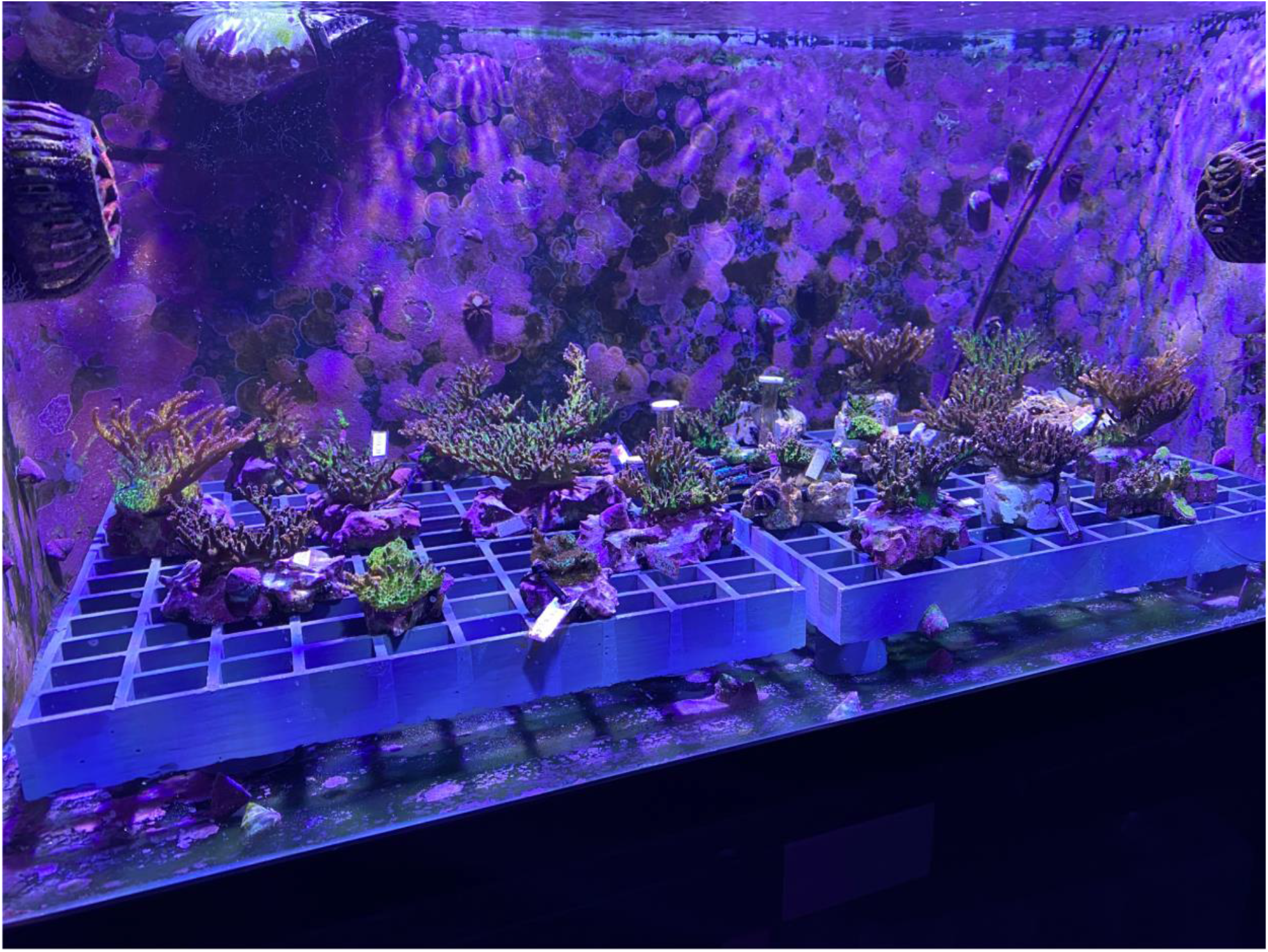
Photo of the F1s produced in 2019. This photo was taken shortly after tissue samples were collected from these individuals, in June 2021 when the F1s were two years old.

### Identity by Descent

We determined kinship among the corals in the system with an identity-by-descent approach. Imported spawners collected from the wild had low identity-by-descent probabilities (pi_hat<0.15), indicating that we collected unrelated individuals as the founders in the system (Supplementary Table 1). Identity-by-descent of parent-parent-offspring trios was consistent with spawn dates of parent corals (Table 1; Figure 2). There was no instance where an offspring appeared to have two parents that spawned on different days.

**Figure 2.**
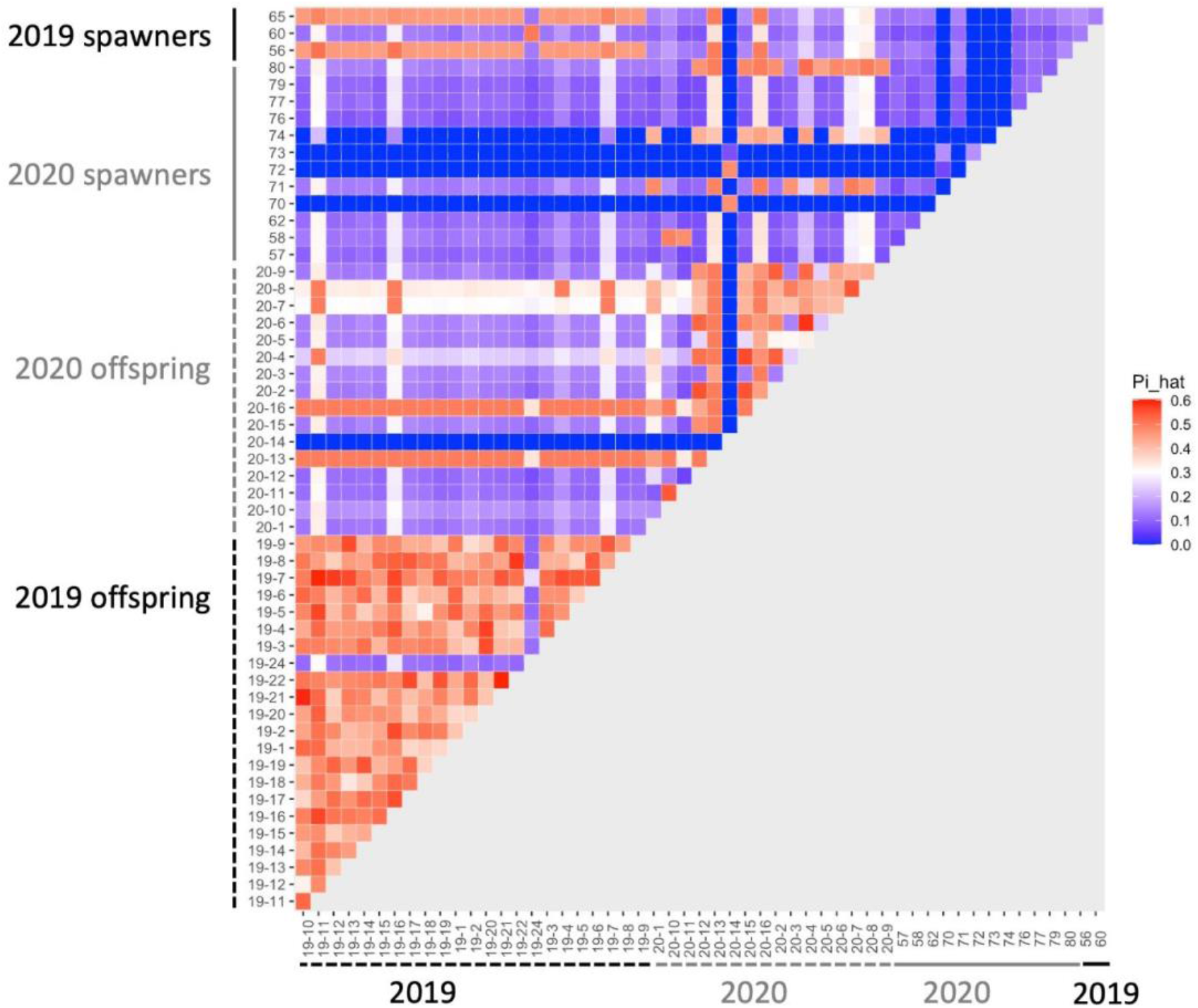
Heatmap of identity-by-descent (IBD) probabilities (pi_hat). Each square shows the probability of IBD for each pair of corals. Theoretical values of IBD indicate that values of ~1 indicates clones, ~0.5 indicates full siblings or parent-offspring, and ~0.25 indicates half siblings.

Based on information from the pi_hat scores we were able to construct pedigrees (Figure 3). Two of the four colonies that spawned in 2019 parented 22 out of the 23 offspring that survived to two years old, meaning that all 22 of these offspring are full siblings (Figure 3a). The other two 2019 spawners produced just one offspring (CA2019-24) that survived to be two years old. CA2019-24 appears to have been parented by CA60 and CA67. We were not able to sequence CA67, but it spawned on the same day as CA60 and CA2019-24 did not have high pi_hat scores with any other individual but CA60, so we infer that CA67 was the second parent.

**Figure 3.**
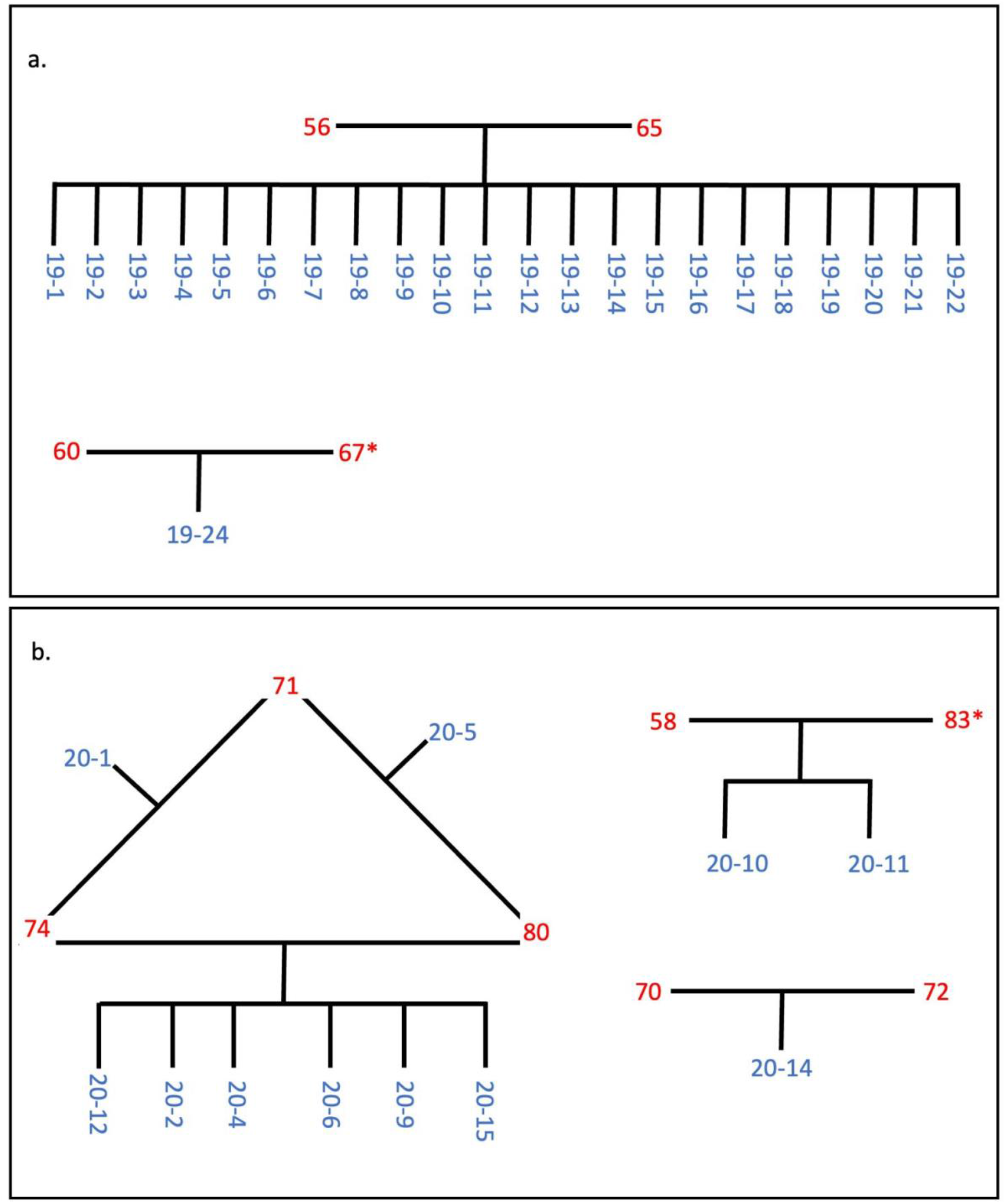
Inferred pedigrees of the parents (red) and offspring (blue) for a.) the 2019 cohort and b.) the 2020 cohort, based on identity-by-descent probabilities (Supplementary Table 2) and spawn dates (Table 1). Colonies with an asterisk were not sequenced, but were inferred to be the parent based on their spawn date and an offspring’s lack of IBD probability of ~0.5 to more than one parent.

For the 2020 spawn, CA74 and CA80 produced six offspring together, and CA74 and CA80 each produced one additional offspring with CA71 (Figure 3b). These two additional offspring, CA2020-1 and CA2020-5, are half siblings of the other four offspring. In addition, CA2020-10 and CA20-11 appear to have been parented by CA58 and CA83, and CA70 and CA72 parented CA20-14. We were not able to sequence CA83, but it spawned on the same day as CA58, and CA2020-10 and CA20-11 did not have high pi_hat scores with any other individual but CA58, so we infer that CA83 must have been the second parent.

There were four offspring from 2020 that had improbably high pi_hat values (i.e., those associated with half- or full siblingship) with nearly every other coral in the dataset, including the 2019 offspring (Figure 2). To check that this was not a result of human error or contamination, we re-extracted DNA from CA20-7 and CA20-8, and sequenced an additional high-coverage full genome library for each of these samples. The new libraries also yielded extremely high pi_hat values with the other corals. These four offspring also display much higher heterozygosity than expected, or than observed in the other corals sequenced (mean percent of sites that were heterozygous in an individual across all samples: 22.4 +/- 5.7%; percent heterozygous for the four corals with IBD scores: CA2020-7: 27.3%; CA2020-8: 28.8%; CA2020-13: 30.1%; CA2020-16: 30.2%).

### F_ST_ outliers

We identified SNPs where the aquarium-bred offspring population significantly deviated from the wild-sourced spawning population by calculating F_ST_ as well as difference in allele frequencies between the spawners and the F1s at each SNP for each cohort. Across all 12,994,408 SNPs, the mean F_ST_ between the 2019 spawners and offspring is 0.02 ± 0.28 (2 s.d.) and the mean F_ST_ between the 2020 spawners and offspring is 0.01 ± 0.11 (2 s.d.) (Figure 4). Of these SNPs, 88,856 were at or above the 99th percentile of F_ST_ values for the 2019 cohort (F_ST_ ≥ 0.47), and 121,419 were at or above the 99th percentile (F_ST_ ≥ 0.20) for the 2020 cohort (Figure 5a,b), and therefore considered outlier SNPs. Of the outlier SNPs for each cohort, 1,442 were shared between both cohorts (Figure 5c). Of the 1,442 SNPs that were outliers in both cohorts, 887 showed a shift in allele frequency in the same direction in both cohorts, where alternate allele frequency either increased from spawners to offspring in both years or decreased from spawners to offspring in both years (Figure 5c). We designated these 887 SNPs as candidate loci potentially undergoing selection in our captive-bred and lab-reared coral. Across all SNPs the mean absolute value change in allele frequency from parents to offspring was 0.06 ± 0.15 (2 s.d.) in the 2019 cohort and 0.08 ± 0.14 in the 2020 cohort (Figure 5d). By comparison, for the 887 shared outlier SNPs, the mean absolute value change in allele frequency from parents to offspring was 0.32 ± 0.09 (2 s.d.) in the 2019 cohort and 0.29 ± 0.13 in the 2020 cohort.

**Figure 4.**
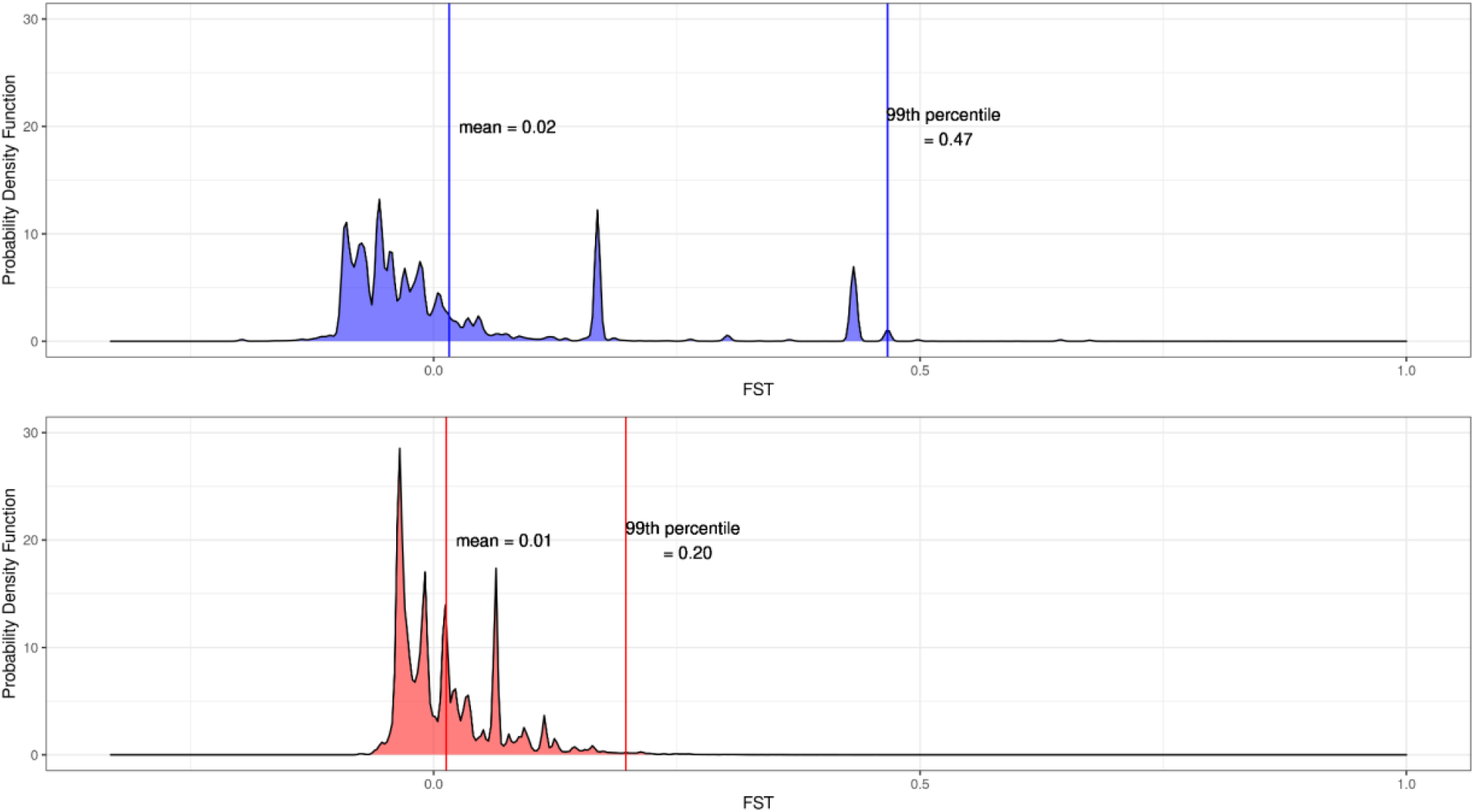
Distributions of F_ST_ at 12.9 million SNPs between spawners and offspring for a.) the 2019 cohort and b.) the 2020 cohort. Vertical lines indicate values of the mean and the 99th percentile for each cohort.

**Figure 5.**
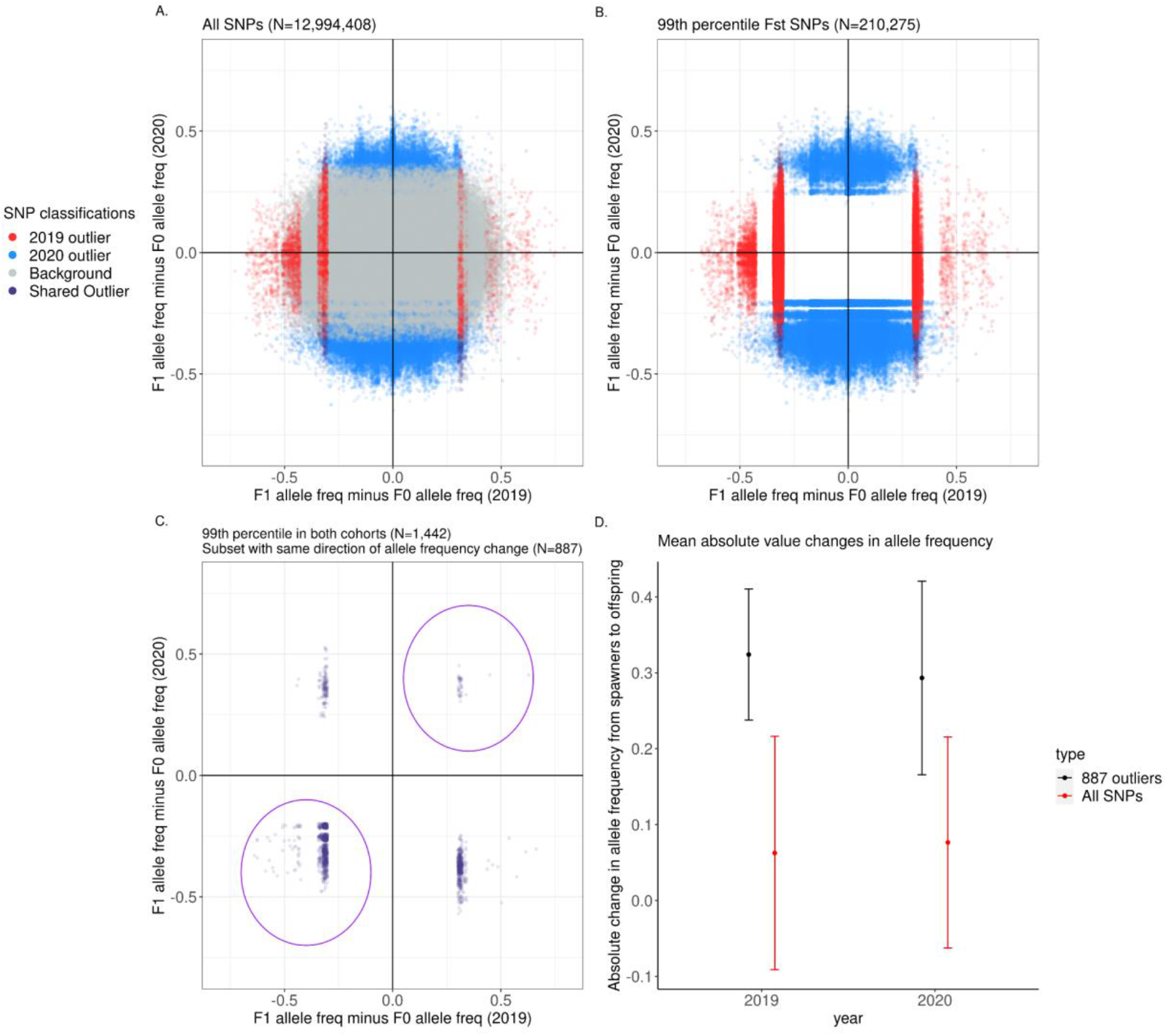
The allele frequency difference between the F1 and F0 in 2019 compared to that in 2020. Gray dots are SNPs that fell outside the 99th percentile of F_ST_ values of all SNPs, red dots are SNPs that fell in the 99th percentile in the 2019 cohort alone, blue dots are SNPs that fell in the 99th percentile in the 2020 cohort alone, and purple dots are SNPs that fell in the 99th percentile in both cohorts. A.) Displays all SNPs in the dataset, B.) displays the 99th percentile SNPs in the dataset, and C.) displays the 99th percentile SNPs that were shared between the 2019 and 2020 cohorts. The purple dots circled in the first and third quadrants in C.) are the 887 outlier SNPs used in the rest of the analyses. D.) Comparison of the mean absolute change in allele frequency for all SNPs (red) and for the 887 outliers (black) between the spawners and the surviving offspring in each year. The dots represent the mean, with the vertical bars representing ± 2 s.d.

### Inheritance in the multi-sibling families

We compared the allele frequency of the 887 outlier SNPs with the allele frequencies of all SNPs for both the 2019 22-sibling family and the 2020 6-sibling family when the alternate allele frequency of the two parents was equal to 0.25, 0.5, and 0.75 (i.e., when the two parents did not share the same homozygous genotype at a SNP; Figure 6). Under Mendelian inheritance, if there is no selection acting on a particular allele, then the allele frequency in the offspring is expected to approximately equal the allele frequency of their parents. For instance, if the parent genotypes are *AA* and *Aa*, the allele frequency of *a* is 0.25, and the expected genotype ratios of the offspring of these two parents would be ½ *AA* and ½ *Aa*, resulting in an expected offspring allele frequency of *a* equal to 0.25.

**Figure 6.**
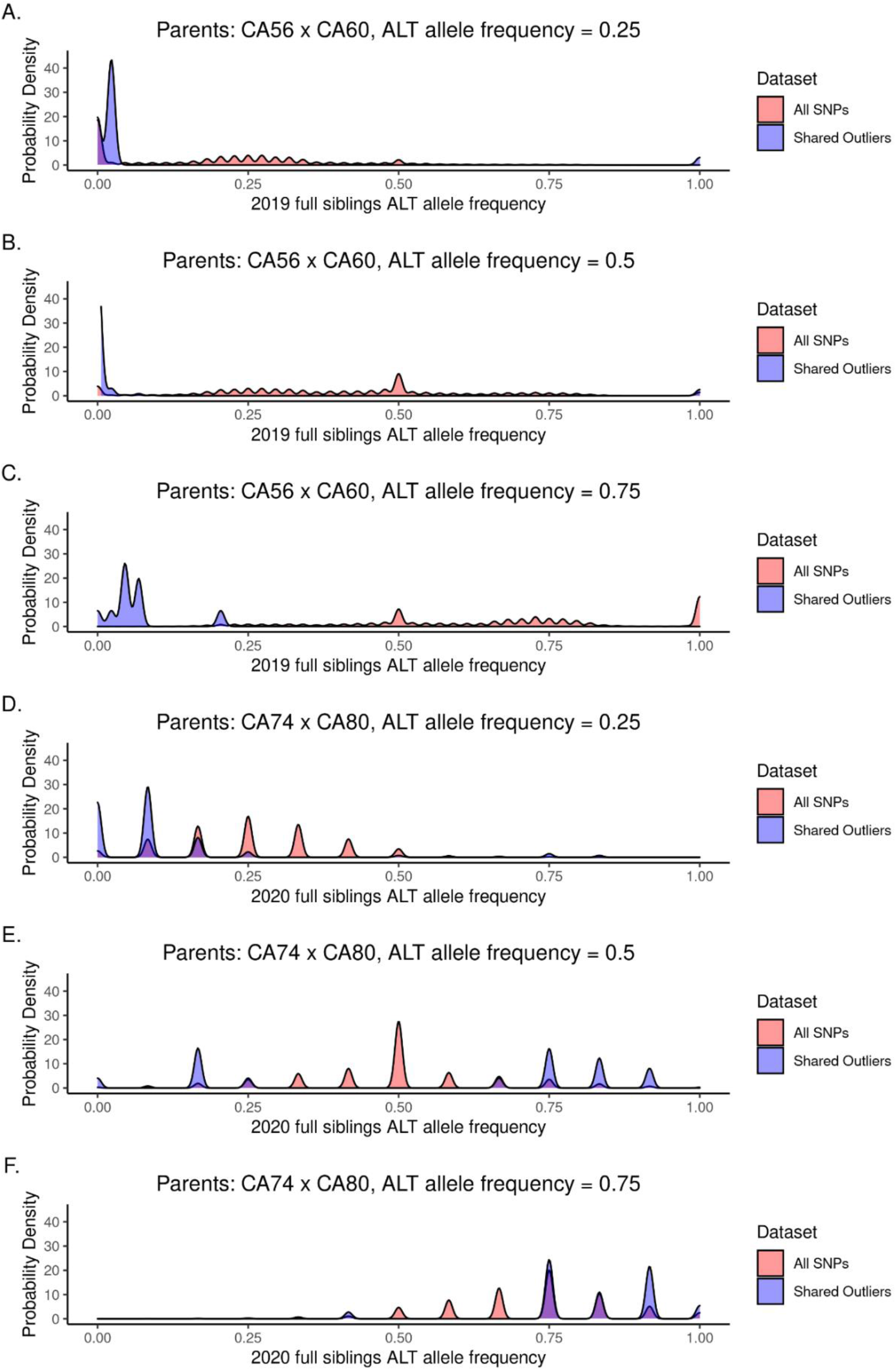
The probability density of allele frequencies for the 22 full siblings produced in 2019 (A-C) and the 6 full siblings produced in 2020 (D-F). The probability densities are shown for the SNPs where the two parents had alternate allele frequency of 0.25 (A and D), 0.5 (B and E), and 0.75 (C and F).

We tested the goodness-of-fit for alternate and reference allele counts for each SNP where the expected offspring alternate allele frequency was 0.25, 0.5, or 0.75, based on the parent alternate allele frequency. Cases where both parents have the same homozygous genotype for either the reference or alternate allele result in parent alternate allele frequency of 0 or 1, so these sites were disregarded for these analyses. P-values for the goodness-of-fit χ^2^ test at each SNP are reported in Supplementary Tables 3 and 4. Across all SNPs where parent alternate allele frequency was equal to 0.25, 0.5, or 0.75 (5,106,603 SNPs for the 2019 family and 4,596,732 SNPs for the 2020 family), 55.6 % are significantly different (p < 0.05) from the expected allele frequency in the 2019 siblings and 10.8% are significantly different from the expected allele frequency in the 2020 siblings. Across the shared outlier SNPs where parent allele frequency was equal to 0.25, 0.5, or 0.75 (873 SNPs for the 2019 family and 129 SNPs for the 2020 family), 100% are significantly different from the expected allele frequency in the 2019 siblings and 37.2% are significantly different from the expected allele frequency in the 2020 siblings.

In the 2019 siblings, for SNPs where the parent allele frequency is 0.25 and 0.5, the offspring allele frequencies are bimodally distributed around 0 and 1, indicating little heterozygosity in offspring at these sites (Figure 6a,b). When the parent allele frequency is 0.75, the offspring allele frequency is bimodally distributed around 0 and 0.25 (Figure 6c). For the six full siblings produced in 2020, the 887 outlier SNPs fit expected allele frequency distributions more closely (Figure 6d-f).

### Hardy-Weinberg Equilibrium

To test whether the aquarium corals fit Hardy-Weinberg equilibrium expectations, we compared the alternate allele frequency with genotype frequency for the three possible genotypes (homozygous reference, heterozygous, and homozygous alternate) at all SNPs and the 887 shared outlier SNPs (Figure 7). We calculated Hardy-Weinberg equilibrium exact test statistics using plink2 to identify SNPs whose observed heterozygosity was significantly different from what would be expected under Hardy-Weinberg equilibrium (Supplementary Tables 5-8). Under Hardy-Weinberg equilibrium, the relationships between allele frequency and genotype frequency are expected to be:

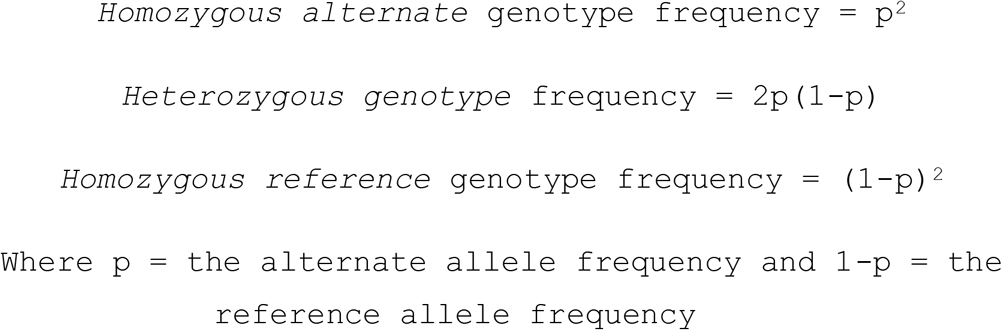

**Figure 7.**
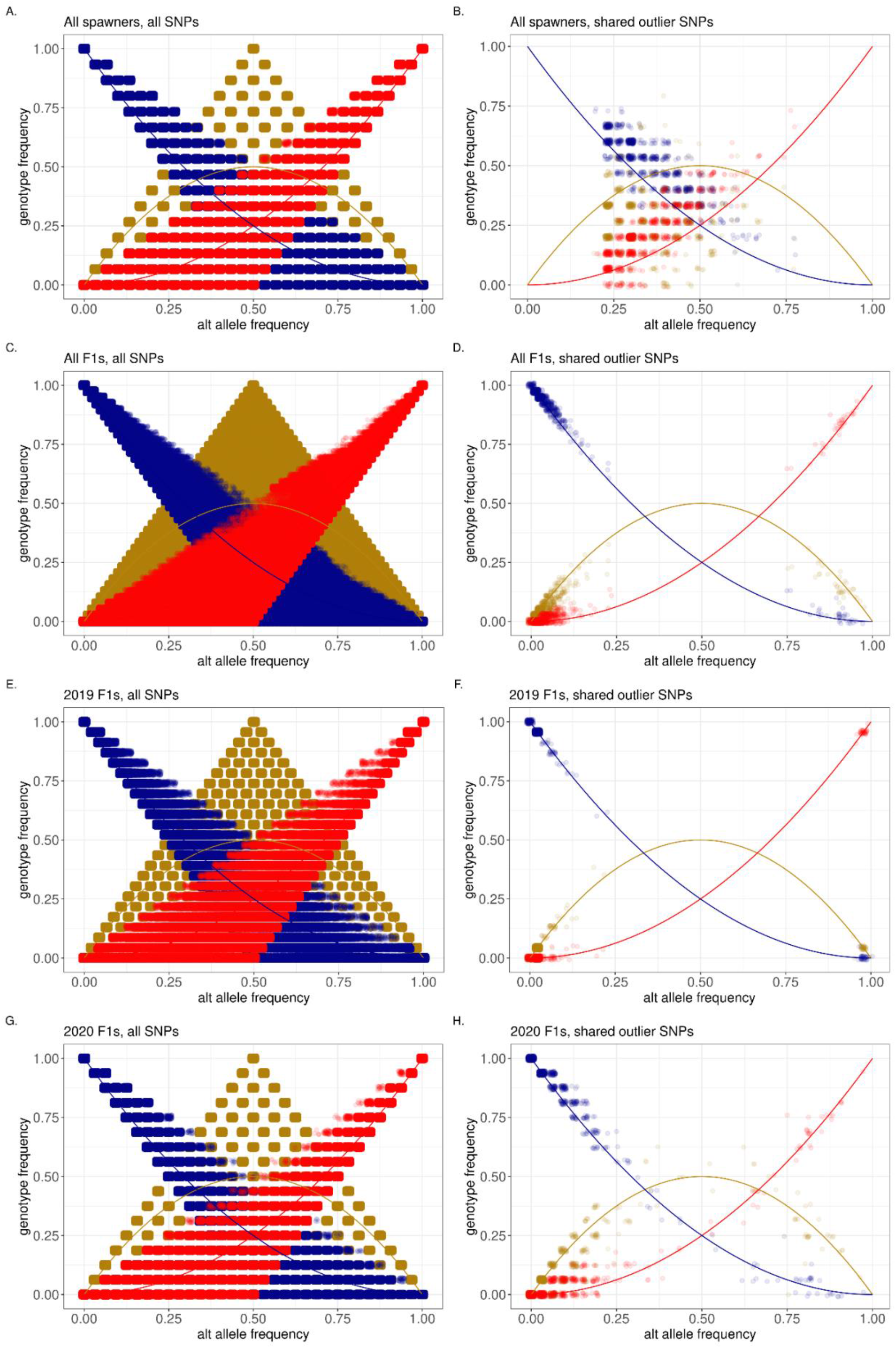
Relationships between the alternate allele frequency and genotype frequency for three genotypes: homozygous reference (blue), heterozygous (gold), and homozygous alternate (red). Equations describing the relationship between alternate allele frequency and genotype frequency expected under Hardy-Weinberg equilibrium for each genotype are represented by lines, while the points are the observed values seen in individual SNPS in each dataset. The observed relationships are shown for a.) all spawners at all SNPs, b.) all spawners at the 887 shared outlier SNPs, c.) all F1s at all SNPs, d.) all F1s at the shared outlier SNPs, e.) the 2019 F1s at all SNPs, f.) the 2019 F1s at the 887 outliers, g.) the 2020 F1s at all SNPs, and h.) the 2020 F1s at the 887 outliers.

For the complete set of wild-sourced spawners, across all 12,994,408 SNPs, 4.3% had a significantly different observed heterozygosity (p < 0.05) than would be expected under Hardy-Weinberg equilibrium (Figure 7a). In comparison, 20.2% of the 887 shared outlier SNPs had a significantly different observed heterozygosity than would be expected under Hardy-Weinberg equilibrium (Figure 7b). Across all F1s produced in 2019 and 2020, across all SNPs, 13.1% had a significantly different observed heterozygosity than expected under Hardy-Weinberg equilibrium (Figure 7c) compared to 8.9% in the shared outliers (Figure 7d). Across the 23 F1s produced in 2019, across all SNPs, 8.2% have heterozygosity that violates Hardy-Weinberg equilibrium (Figure 7e). Most of the shared outliers are at or near fixation, however, just one of the 887 SNPs (0.001%) has a heterozygosity that violates Hardy-Weinberg equilibrium (Figure 7f). Across all SNPs for the 2020 F1s, 5.7% violate Hardy-Weinberg equilibrium (Figure 7g), and 8.1% of the shared outliers violate Hardy-Weinberg equilibrium (Figure 7h).

### GO enrichment of outliers

To explore gene networks that may have been under selection across cohorts, we tested for an enrichment of GO biological functions and pathways in the set of shared outlier SNPs. A total of 42 molecular function and 189 biological pathway GO terms were enriched in the set of genes associated with the shared outlier SNPs (p < 0.05; see full list in Supplementary Table 7). Notably, the functions syntaxin/SNARE binding and GTPase activator/regulator activity and several vesicle-mediated transport pathways were among the top enriched terms (Supplementary Table 7).

## DISCUSSION

The 2019 and 2020 coral spawns at California Academy of Sciences were among the first in the United States to successfully produce aquarium-born offspring that have survived to over three years old (at the time of this publication). Unlike clonal propagation, successful sexual reproduction allows corals to maintain standing genetic variation and produce new genotypes through recombination. This genetic diversity will be essential for coral populations’ capacity to adapt to environmental change. By sequencing the spawners as well as the offspring that lived to be at least one year old, we are able to describe the genome-wide variation of aquarium-bred corals for the first time. With this information, we are equipped to make recommendations about how to maximize genetic variation and minimize the effects of genetic drift and adaptation to the aquarium environment in future coral breeding efforts.

### Spawning and juvenile coral rearing methods

Given the importance of a genetically diverse broodstock, captive breeding programs should maximize the number of genetically distinct individuals that synchronously spawn on the same day and time (as opposed to segmented spawning, where individuals spawn on consecutive days) to produce the most heterogeneous starting population possible. Further work is needed to develop spawning cues that operate on fine scales (i.e., days, hours, or minutes) to facilitate the most synchronous spawns possible. Higher synchronicity in spawning of *Acropora hyacinthus* colonies has been achieved in an aquarium than was observed in this study (Craggs et al., 2018). Interestingly, ex situ corals have been observed to spawn a few days after their in situ counterparts across a variety of species, including Pacific corals and Caribbean corals (Craggs et al., 2017; Neely et al., 2020). To inform best practices for restoration fertilization, genetic heterogeneity of embryos yielded from batch fertilization (as in this study) versus controlled crosses of pairs of individuals (e.g Humanes et al., 2021) should be compared to determine the method that maximizes genetic variation. The development of standard husbandry protocols for grow-out may also help optimize genetic diversity of broodstock by maximizing survivorship, and therefore minimizing bottlenecks, at each life stage. This may include species-specific protocols for settlement, symbiont inoculations, feeding regimes, and cleaning/grazing regimens (Levenstein et al., 2021; O’Neil et al., 2021; Rahnke et al., 2022).

### Parentage of aquarium-bred corals

Differential parentage is apparent in our system. In the 2019 cohort, 22 out of 23 offspring that survived to be two years old are full siblings that share the same two parents. Skewed contributions of spawners to surviving offspring have also been found in other conservation breeding programs, including perch (Attard et al., 2016) and another coral species, *Acropora palmata* (Hagedorn et al., 2021). The 2020 cohort represented a more even contribution of spawner genotypes. The eleven offspring for which parents could be identified came from seven of the thirteen colonies that spawned. There were also four 2020 offspring for whom parents could not be readily assigned, and which showed implausibly high pi_hat values with nearly all other spawners and offspring in the dataset, as well as anomalously high heterozygosity across the genome. This suggests that these are chimeras, or colonies made up of more than one sexually produced genotype. While the prevalence of chimerism in wild *Acropora hyacinthus* is only 3% (Schweinsberg et al., 2015), it is likely that chimerism occurs more frequently in aquarium-bred corals due to limited dispersal and settlement area (Puill-Stephan et al., 2012). Some studies indicate that chimeras may be more resilient to thermal stress (Huffmyer et al., 2021) and disease (Williamson et al., 2022), so their increased prevalence in aquarium populations may be adaptive.

### Allele frequency comparisons between spawners and offspring

Allele frequencies did not differ significantly between spawners and offspring across the full genome. This is likely because one generation and a small population size is not sufficient to observe drastic changes, particularly as the starting population was highly heterozygous. However, there was a subset of SNPs for which allele frequency differences were anomalously high in both the 2019 and 2020 cohorts. Follow-up studies in other systems, and in later generations if the current F1s later produce F2s, may help to indicate their importance, or lack thereof. So far just one F2 generation has been produced in an aquarium (Craggs et al., 2020), but with expanding resources and knowledge in coral breeding and husbandry, this will soon become more common and allow for further study of allele frequency changes over generations.

### Mendelian inheritance anomalies in the outliers SNPs

It is not clear from our data which phenomenon is the largest contributor to differential parentage in the surviving F1 corals. Possible causes include certain spawners producing a disproportionate quantity of gametes in the batch fertilization pool, certain spawners producing more vigorous sperm, certain spawners being more genetically compatible with others in the population, or differential survival of certain genotypes at early (embryonic, larval, young recruit) life stages. The strongest evidence that this bias is at least in part explained by differential survival of certain genotypes comes from the deviation from Mendelian expectations in the outlier SNPs for the 22 full siblings from two highly successful 2019 parents (Figure 5a-c) and the 6 full siblings produced in 2020 (Figure 5d-f). If there was no selection acting on the shared outlier SNPs, then the allele frequency of the F1s in both families should have matched that of their parents. Instead, we see fixation for one allele among full siblings at most of the shared outlier SNPs. 100% and 37.2% of shared outlier SNPs deviate significantly from the allele frequency expected given their parents’ allele frequency at that SNP in the 2019 and 2020 full siblings, respectively. This is more consistent with directional selection than with genetic drift, as it is unlikely that alleles would fix or become nearly fixed at the same locus for two independent sets of siblings spawned in two different years from different parents. In contrast, the SNPs that fell in the 99th percentile of F_ST_ values in just one cohort or the other (i.e., the red and blue points in figures 4a and 4b) are more likely to be outliers due to genetic drift, because the stochastic nature of drift makes it unlikely to act on the same SNPs in two independent populations.

This phenomenon, in which certain sites become fixed or nearly fixed in a single generation while the majority of SNPs genome-wide maintain expected levels of heterozygosity, may be unique to broadcast spawners and other animals that produce many offspring each time they reproduce. Each spawning event results in millions of embryos, and that embryo pool is highly heterogenous (Kitanobo et al., 2022). In the wild and in the aquarium, the vast majority of coral embryos do not survive to maturity, so genetic drift and selection during the earliest life stages may be very strong. One way to test this hypothesis would be to model the starting heterogeneity found in an initial embryo pool, then simulating different outcomes by changing stochastic drift and selection coefficient parameters in the model.

### Hardy-Weinberg equilibrium in aquarium-bred corals

Hardy-Weinberg equilibrium is a mathematical description of the expected relationships between allele and genotype frequency in a population in the absence of migration, mutation, selection, and assortative mating (Hardy 1908; Weinberg 1908). When populations deviate from Hardy-Weinberg equilibrium, it can indicate that inbreeding, population stratification, or other evolutionary forces are acting on the population (Wigginton et al., 2005). We tested how many sites across the genome had a significantly different heterozygosity than would be expected under Hardy-Weinberg equilibrium for the spawner and offspring populations. Overall, the vast majority of sites in both the wild and aquarium-raised corals were in Hardy-Weinberg equilibrium, and the F1 cohorts maintained high heterozygosity across the genome. Within the 887 shared outlier SNPs, the 2019 F1s show much lower heterozygosity and higher fixation than their parents, while the 2020 F1s maintain much higher heterozygosity. The difference between the two cohorts is likely due to the fact that all but one of the 2019 offspring came from the same two parents, whereas there was a higher diversity of parentage in the 2020 offspring.

The fact that the 2020 F1s are still largely in Hardy-Weinberg equilibrium and maintain high heterozygosity, even at the 887 outlier SNPs indicates that, for corals that are highly heterozygous to begin with, just seven successful parents can produce a genetically diverse set of aquarium-bred offspring. In addition, the minimal increase in the number of SNPs that deviate from Hardy-Weinberg expectations may suggest that overall there has not been a great deal of selection or allele frequency changes due to other factors in one generation. This may bode well for out planting aquarium-bred corals into the wild, and indeed it has been shown that aquarium-bred corals can survive and grow successfully in the wild (Henry et al., 2021), in contrast with studies that have demonstrated fitness and phenotype changes in hatchery-raised fish within one generation (Kostow, 2004; Wakiya et al., 2022).

### Genes affected by the shared outlier SNPs

Though we are cautious not to overinterpret our Gene Ontology enrichment results (see Methods for limitations), we highlight the most prominent enriched functions here. We do not suggest that these functions are definitively under selection in aquaria, but SNPs related to these functions were significant F_st_ outliers in the aquarium-bred offspring, and therefore merit further consideration and study to determine whether the aquarium environment affects these functions in a manner that is different from what juvenile corals would experience in the wild. Several vesicle transport, and particularly exocytosis, functions are enriched in the shared outliers, suggesting that genes involved in expelling vesicle contents may be important for the success of aquarium-bred corals. Syntaxin binding with synaptotagmin (a gene that lies directly upstream of a shared outlier SNP) is well established as a critical activator of vesicle exocytosis; transmembrane transport of vesicles occurs when an influx of calcium ions bind to synaptotagmin-syntaxin complexes (Jena, 2009; Ohya et al., 2009). Previous work has demonstrated that synaptotagmin-like protein is activated during light-induced bleaching in *A. microphthalma* (Starcevic et al., 2010). Additionally, Rab GTPases regulate membrane and vesicle transport (Deneka et al., 2003) and have been shown to play a key role in the establishment and maintenance of endosymbiosis in the model anemone *Exaiptasia* (Chen et al., 2003). Given that endosymbionts are encapsulated in vesicles within coral host cells, one possibility is that selection may be acting on vesicle transport as it relates to endosymbiont uptake and expulsion. An exciting future direction that may provide additional clues is to determine when in development outlier-associated genes are relevant, which could be evaluated with gene and protein expression data of larvae across developmental stages. Future research is required to elucidate how and why the coral-algal relationship may be different in the lab environment than in the wild environment.

### Recommendations

In any conservation breeding program, maximizing genetic variation and minimizing detrimental effects due to inbreeding or selection for the zoo or aquarium environment are crucial for facilitating good outcomes once the conservation-bred individuals are released into the wild (Frankham, 2008; Lacy et al., 2018). Based on our results from the 2019 and 2020 spawns at California Academy of Sciences, we recommend the following guidelines for maximizing standing genetic variation in an aquarium-bred coral population.

1. Choose a highly heterozygous breeding stock if possible.
2. Start with at least 10 individuals in the breeding stock-not all will spawn, and often they will not spawn on the same night. Aiming for at least seven spawners that give rise to surviving offspring can ensure a F1 population that is highly heterozygous and in Hardy-Weinberg equilibrium (Figure 7g, h).
3. Equalize the size of families, so that no one or two parents is excessively successful compared to the others. One way to control this in corals and other broadcast spawners is to do fertilization via reciprocal crosses rather than batch fertilization, using known volumes of sperm at equal concentrations.
4. Maximize survivorship at every step-through symbiont inoculations, introduction of grazing invertebrates like snails and urchins, etc.- to reduce bottleneck effects and consequent genetic drift at certain life stages.
5. Introduce aquarium-bred F1s back out onto the reef each generation, rather than keeping aquarium-bred lines going for multiple generations, to reduce the effects of lab selection.
6. Introduce a few new wild breeders each generation, as in Sahashi and Morita (2022), preferably in the form of cryopreserved sperm so that colonies do not have to be taken off of the reef each year (as in Hagedorn et al., 2021; Howell et al., 2021).

### Next steps

There are still many unresolved questions about best practices in breeding corals for conservation purposes, especially with the express aim of maximizing standing genetic variation in aquarium-bred offspring. Future research to determine which life stage is most subject to selection pressures will be crucial to our understanding of where the most resources for maximizing survivorship should go. Monitoring allele frequency changes throughout the first few days of embryo and larval development may elucidate where the major bottlenecks occur, and which genetic variants are most detrimental and beneficial in getting a given embryo through to the juvenile coral colony life stage.

Another avenue of research, one that may come at odds with the principles of maximizing standing genetic variation, is to experimentally select for desired traits in aquarium-bred corals. There is a lot of appeal in assisted evolution, or artificially selecting for animals that are best at surviving higher temperature, lower pH, or other environmental factors predicted to change in the ocean in the coming decades. Whether this can be done in a way that does not also eliminate too much variation across the genome at other loci unrelated to a given trait of interest remains to be tested.

Further, our results hint that specific genes and biological pathways may be under selection in the aquarium environment. The implication of this is two-fold; it may be possible to predict genetic variants and associated traits that underlie embryo and larval success in the aquarium, and genetic variants selected for in the aquarium may or may not be well-suited for success in the wild. In both cases it will be beneficial to evaluate functional traits in lab strains, such as response to environmental stressors and characteristics of endosymbiont uptake.

## ACKNOWLEDGEMENTS

This study was supported by the California Academy of Sciences’ Hope for Reefs Initiative, with generous support from Carnegie Science, the Kingfisher Foundation, Autodesk, and the Hearst Foundation. The following companies donated aquarium equipment and other products: Neptune Systems, Ecotech Marine, Two Little Fishies, and Coral Polyp Lab.

## AUTHOR CONTRIBUTIONS

Study conception and design: EHLN and RA; animal care, system design, and resources: LL, JCD, RR, KL, RS, SY, FD, LK, and RA; data collection: EHLN and RA; analysis and interpretation of results: EHLN and CYP; draft manuscript preparation: EHLN; supervision: RA. All authors helped shape the research and provided critical feedback on the manuscript.

## CODE AND DATA AVAILABILITY STATEMENT

Raw fastqs are acessioned at NCBI SRA with accession numbers SAMN30634691-30634742. Code used to run analyses described in this paper can be found in the AquariumBreedingGenomics Github repository: https://github.com/eloralopez/AquariumBreedingGenomics

## Appendix 1. Key commands used in analysis

### BLAST outlier-associated A. hyacinthus transcripts against local N. vectensis cDNA database for GO term matchup

blastn -db Nematostella_vectensis.ASM20922v1.cdna -query trulyshared_tx_all.fasta -task blastn -word_size 11 -outfmt 6 -out trulyshared_tx_all.Nv-cdna-blastn.out

### Choose top hit per transcript based on bit score (1st) and e-value (2nd)

sort -k1,1 -k12,12nr -k11,11n trulyshared_tx_all.Nv-cdna-blastn.out | sort -u -k1,1 --merge > top_hits.trulyshared_tx_all.Nv-cdna-blastn.out

